# Features of Functional Human Genes

**DOI:** 10.1101/2020.10.10.334193

**Authors:** Helena B. Cooper, Paul P. Gardner

## Abstract

Proteins and non-coding RNAs are functional products of the genome that carry out the bulk of crucial cellular processes. With recent technological advances, researchers can sequence genomes in the thousands as well as probe for specific genomic activities of multiple species and conditions. These studies have identified thousands of potential proteins, RNAs and associated activities, however there are conflicting conclusions on the functional implications depending upon the burden of evidence researchers use, leading to diverse interpretations of which regions of the genome are “functional”. Here we investigate the association between gene functionality and genomic features, by comparing established functional protein-coding and non-coding genes to non-genic regions of the genome. We find that the strongest and most consistent association between functional genes and any genomic feature is evolutionary conservation and transcriptional activity. Other strongly associated features include sequence alignment statistics, such as maximum between-site covariation. We have also identified some concerns with 1,000 Genomes Project and Genome Aggregation Database SNP densities, as short non-coding RNAs tend to have greater than expected SNP densities. Our results demonstrate the importance of evolutionary conservation and transcription for sequence functionality, which should both be taken into consideration when differentiating between functional sequences and noise.

## Introduction

The definition of “functional” when it comes to genes is extensively debated, with the most contention being whether biochemical activity alone, or evidence of evolutionary selection, is required to conclude whether a gene is functional (Eddy 2012; Doolittle and Brunet 2017; Graur 2017; ENCODE Project Consortium 2012; ENCODE Project Consortium et al. 2020; Cheetham et al. 2020; Godfrey-Smith 1994). From the perspective of selection, between 3% and 15% of the human genome is considered to be under purifying selection based upon comparisons of homologous regions in eutherian genomes (Ohno 1972; Siepel et al. 2005; Lunter et al. 2006; Davydov et al. 2010; Ponting and Hardison 2011; Lindblad-Toh et al. 2011; Rands et al. 2014; Graur 2017). Therefore, from an evolutionary selection perspective, these regions are very likely to be functional. However, evolutionary estimates may be negatively influenced by issues with the accuracy of nucleotide alignments (Gardner et al. 2005; Freyhult et al. 2007) and the calibration of evolutionary models (Pheasant and Mattick 2007). The evolutionarily constrained proportion of the genome is considerably smaller than the reports summarising large-scale screens for biochemical activity from projects such as ENCODE (ENCODE Project Consortium 2012; ENCODE Project Consortium et al. 2020; Pheasant and Mattick 2007). ENCODE authors concluded that around 80% of the human genome can be assigned a “biochemical function”, which meant a sequence could be defined as functional so long as it participated in at least one biochemical event, such as transcription or a regulatory process, in one or more cell types (ENCODE Project Consortium 2012). This conclusion is controversial with researchers familiar with the mechanisms of genome size variation, and an awareness of biological and technical noise, who have argued that genomic activity could be attributed to biological or experimental noise, rather than function (Germain et al. 2014; Eddy 2012; Doolittle 2013; Doolittle et al. 2014; Doolittle and Brunet 2017).

The aim of our study is to evaluate the relative importance of six groups of genomic features, which were selected based on their potential to predict gene functionality, by comparing human protein-coding and non-coding RNA (ncRNA) genes to non-genic control regions. These are intrinsic sequence features, sequence conservation, transcription, genomic repeat association, protein-coding or RNA selection features and population variation. We have ranked features that are strongly predictive of assigned gene functionality, using both a Spearman correlation analysis and random forest classification models. The results of both approaches are largely concordant, ranking measures of inter-species evolutionary and transcription highly, while selection measures based upon population variation data had low rankings. In summary, we find that sequence conservation, followed by evidence of transcription, were the most associated with assigned sequence functionality.

## Results

Sequences that are assigned as “functional” in this study are annotated protein-coding, short ncRNA and long ncRNA (lncRNA) genes from the HUGO Gene Nomenclature Committee (HGNC) database, and are therefore linked to documented evidence of function (Braschi et al. 2019). Chromosome Y and mitochondrial DNA are excluded due to excess low complexity sequence for the former and different selection pressures for the latter (Quintana-Murci and Fellous 2001). Both ncRNA datasets excluded ribosomal RNAs (rRNA) due to their large sequence length, unusual expression and sequence conservation patterns and uncertain placement in the reference human genome (Agrawal and Ganley 2018). A maximum of 1,000 sequences were selected at random for each dataset, which are filtered to only include sequences less than 3,000 bp in length to limit length-dependent effects. For protein-coding and lncRNA genes, only exons two and three are included to restrict the analysis to internal exons, thus minimising the possible influence of introns, untranslated regions and promoters. The short ncRNA dataset included small nucleolar RNAs (snoRNAs), transfer RNAs (tRNAs), small nuclear RNAs (snRNAs), microRNAs (miRNAs) and other less represented groups such as guide and Y RNAs.

The ideal negative control sequences for this study are human genomic regions that are guaranteed to be non-functional, while maintaining background levels of sequence content, population variation, conservation, and transcription. In the absence of this ideal, our negative control sequences were generated based on the chromosomal coordinates of each functionally assigned sequence, with the new coordinates being 20 kb upstream and downstream. The 20 kb range was chosen to provide sufficient distance between functional and negative control sequences to prevent genetic linkage influencing the population variation results (Loh et al. 2013). To account for sequence length bias, each negative control sequence had the same sequence length of either the short ncRNA or protein-coding and lncRNA exon it matched with. To ensure that the negative control sequences are not located in known protein-coding genes or ncRNAs, overlaps with annotated genes were removed and the remaining sequences used as negative controls and assigned as being “non-functional”. A fraction of these may have undetermined function, but our expectation is that the bulk of these are non-functional, junk DNA (Palazzo and Gregory 2014).

Features that may be associated with gene functionality were selected from a range of statistics and scores that met the five criteria detailed in the methods (Table S1). Spearman correlation matrices and random forest models were used to assess the strength of associations between genome features and assigned gene function. Protein-coding, short ncRNA and lncRNA sequences are analysed separately, and are only compared to their corresponding group of negative control sequences. Spearman correlation matrices are calculated using Spearman’s Rho, with correlations computed between the assigned functional status (“0” or “1”) and each genome feature. Random forests were generated using a 70% training and 30% test data split, with functionally assigned sequences as “controls” and negative control sequences as “cases”. Variable importance within a random forest was determined using the mean decrease in the Gini-coefficient, which is an estimate of how important a variable is for assigning functionality across all the random trees. The genome features that are associated with function have both a significant Spearman correlation (P<0.05) and a random forest Gini-coefficient above the neutral predictor.

### Random forest model performance

To distinguish between genome features that indicate function and noise in the random forest, we have included a “neutral predictor”, which is the 5’ chromosome coordinate for each sequence. Because the negative control sequences are generated using the assigned functional sequence, the 5’ chromosome coordinate will not differ between the functional and control sequences. Indeed, there is no significant correlation with assigned sequence functionality or a significant difference between protein or lncRNA exons, confirming that this feature is a suitable threshold for non-informative features in a random forest (Figure 1; Table S2).

**Figure 1:**
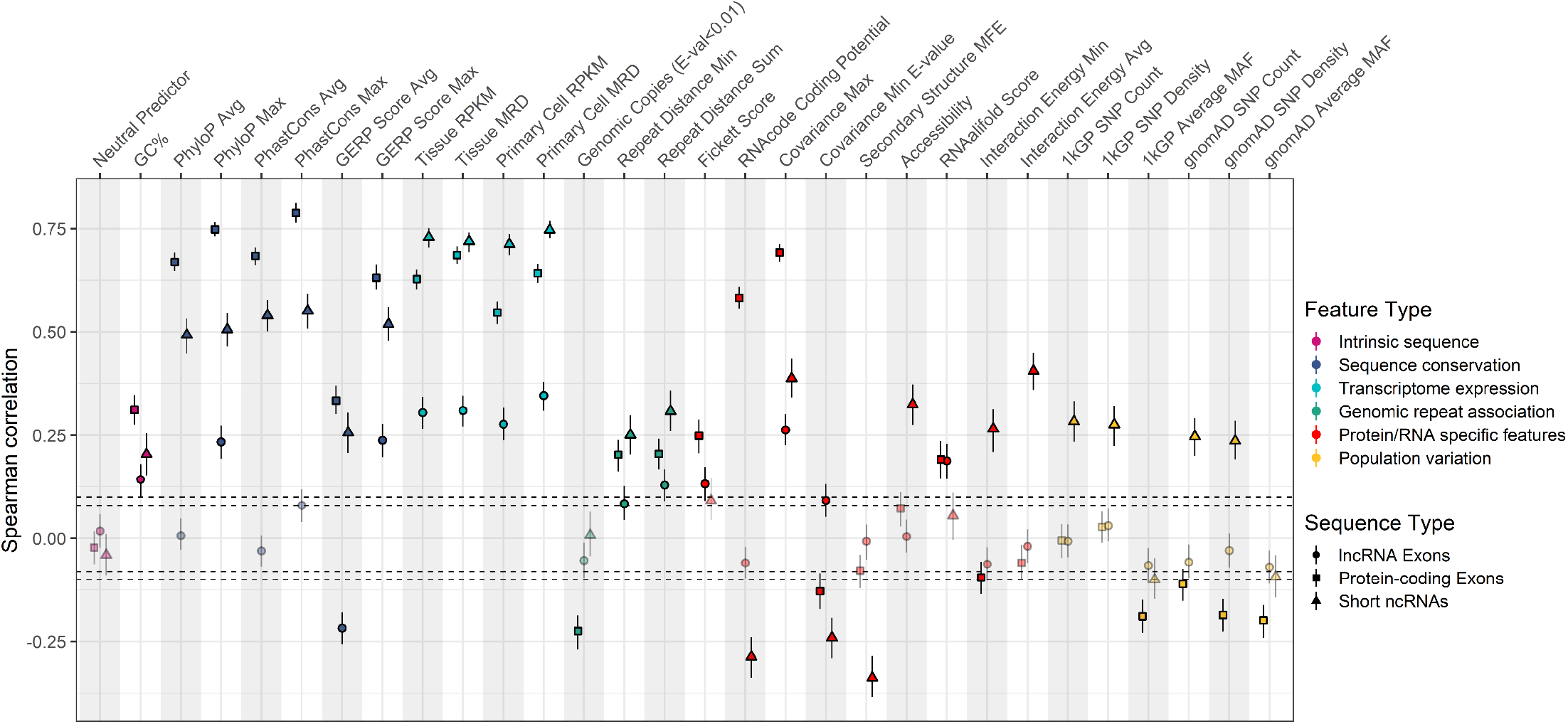
The Spearman correlation between genomic features and the assigned functional status of protein-coding (N^Functional^=1,576 & N^Non-functional^=845), short ncRNA (N^Functional^=803 & N^Non-functional^=635), lncRNA (N^Functional^=1,693 & N^Non-functional^=635) sequences. Each point shows the Spearman correlation of a feature with assigned sequence functionality, which includes the 95% confidence intervals calculated using bootstrapping (replicates=1,000). Non-significant correlations (P>0.05) appear faded, with dotted lines highlighting the threshold of non-significance, which was ±0.094 for protein-coding regions, ±0.110 for short ncRNAs and ±0.082 for lncRNAs. Sequences that had at least one NA recorded for any feature were removed from the Spearman correlation analysis. The full Spearman correlation matrices and values are available in Figures S1-S3.

Prior to identifying features associated with assigned sequence functionality, the classification accuracy for the random forest models are evaluated by calculating the specificity, sensitivity, accuracy, F1 score, Matthew’s correlation coefficient (MCC) and area under the curve (AUC). Overall, the random forest classification models generated for the protein-coding and short ncRNAs sequences perform well based upon our cross-validation results (Table 1). However, the lncRNA classification models are considerably less discriminative, with an average MCC of 0.404 compared to 0.844 and 0.879 for the short ncRNA and protein-coding models respectively (Table 1). These metrics suggest that lncRNA features with a high variable importance are not necessarily suitable predictors of lncRNA functionality, due to the large number of false negatives (Table 1).

**Table 1:** Summary of random-forest performance metrics across assigned functional sequence classification models (Figure 2).

|  | Protein-coding |  |  | Short ncRNA |  |  | lncRNA |  |  |
| --- | --- | --- | --- | --- | --- | --- | --- | --- | --- |
|  | Min | Avg | Max | Min | Avg | Max | Min | Avg | Max |
| Sensitivity | 0.906 | 0.919 | 0.921 | 0.927 | 0.930 | 0.952 | 0.374 | 0.393 | 0.410 |
| Specificity | 0.950 | 0.958 | 0.959 | 0.908 | 0.918 | 0.929 | 0.925 | 0.934 | 0.938 |
| MCC | 0.846 | 0.879 | 0.919 | 0.780 | 0.844 | 0.892 | 0.273 | 0.404 | 0.482 |
| Accuracy | 0.930 | 0.945 | 0.964 | 0.891 | 0.923 | 0.947 | 0.762 | 0.787 | 0.818 |
| F1 Score | 0.898 | 0.921 | 0.947 | 0.882 | 0.912 | 0.940 | 0.398 | 0.504 | 0.581 |
| AUC | 0.973 | 0.982 | 0.991 | 0.968 | 0.980 | 0.991 | 0.722 | 0.792 | 0.835 |

### Feature Analysis

#### Intrinsic sequence features

A high G+C content is often associated with functional protein-coding exons due to its role in exon selection and transcription (Kudla et al. 2006; Amit et al. 2012), but as this trend has also been observed in ncRNAs, this could be a useful feature of functionality (Klein et al. 2002; Haerty and Ponting 2015). For protein-coding and short ncRNA sequences, G+C content is significantly correlated with assigned functionality and its importance ranks above the neutral predictor by the random forests (Figures 1 & 2), with the observed positive correlations following the expected trend. Although these observations apply to the lncRNAs, with G+C importance ranked third by the random forest, this high ranking conflicts with this feature having the fifth lowest Spearman correlation of 0.14 to assigned lncRNA functionality (Figures 2 & S3). Therefore, it is likely the high Gini coefficient reflects the significantly higher G+C content in lncRNA exon two, rather than being representative of the full-length lncRNA, as only exon two produces a significant correlation with lncRNA functionality (Figure S4; Table S2).

**Figure 2:**
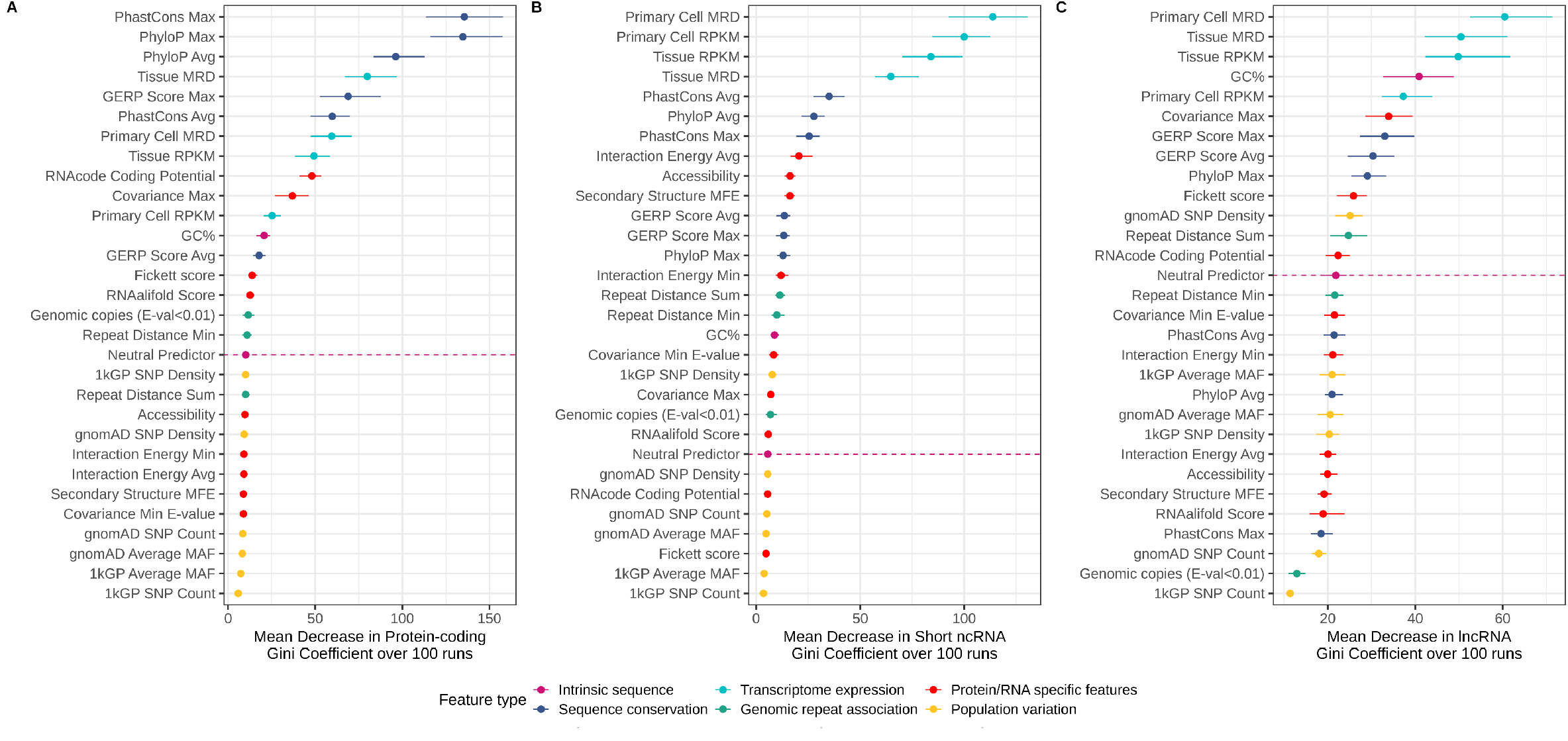
Gini coefficient importance plot from 100 random forest prediction models for assigned protein-coding, short ncRNA and lncRNA sequence functionality. The x-axis is the mean decrease in Gini coefficient averaged across 100 random forest models and the y-axis lists the functionality features, each coloured by feature type. The range of Gini coefficients observed across all 100 classification models is shown for each feature. The neutral predictor, which is the starting chromosome position for each sequence, is labelled with a dotted line. Predictors listed below this line are assumed to not contribute much to the random forest models and are thus unable to differentiate between assigned functional and non-functional sequences. Model performance metrics are available in Table 1. Sequences with NAs recorded had these features approximated by the random forest. **A:** Protein-coding exons (N^Functional^=2,000 & N^Non-functional^=1,104). **B:** Short ncRNAs (N^Functional^=1,000 & N^Non-functional^=895). **C:** lncRNA Exons (N^Functional^=2,000 & N^Non-functional^=790).

#### Sequence conservation features

The presence of conserved genomic regions between species that diverged millions of years ago implies that these are required for survival, and hence are likely to be functional (Doolittle et al. 2014; Doolittle and Brunet 2017). PhyloP, PhastCons and GERP (Genomic Evolutionary Rate Profiling) scores are computed for each nucleotide in the genome based on phylogenetic trees (Pollard et al. 2010; Siepel et al. 2005; Cooper et al. 2005). All sequence conservation features are positively correlated with assigned protein-coding and short ncRNA functionality, indicating that these sequences are generally more conserved than the negative control regions (Figure 1). The conservation features rank well above the neutral predictor by random forest for both sequence types, indicating that sequence conservation is strongly associated with sequence functionality (Figure 2). For the lncRNAs, the only conservation features that have a significant correlation and rank above the neutral predictor are maximum PhyloP and the GERP scores, with the GERP score average producing a negative correlation, suggesting a lack of conservation in lncRNAs (Figures 1 & 2). One possible explanation is that the maximum PhyloP and GERP scores identify conserved functional domains in lncRNA sequences (Figure 1).

#### Transcriptome expression features

If a gene is expressed in at least one tissue during at least one developmental stage, then it is available for selection to act on if there is a functional affect (Kaikkonen and Adelman 2018; Buccitelli and Selbach 2020). We use both the maximum read depth (MRD) and the average number of reads per kilobase of transcript, per million mapped reads (RPKM) to summarise transcription across ENCODE Total and Small RNA-Seq experiments from either 71 human tissue or 140 human primary-cell samples (ENCODE Project Consortium 2012). All transcription features produce significant positive correlations with the functionally assigned sequences, and rank above the neutral predictor in the random forest analysis, implying that transcription is likely associated with gene functionality (Figures 1 & 2). In particular, primary cell MRD is the most informative feature for both ncRNA datasets, whereas sequence conservation is most informative for protein-coding sequences (Figure 2).

#### Genomic repeat associated features

Duplicated genes may be more likely to be functional compared to single-copy genes, suggesting that genomic copy-number could be a potential feature of function (Ohno 1970). To estimate the number of genomic copies per sequence, blastn was used to report significantly similar sequences (E-value<0.01) within the human genome (Altschul et al. 1990). Overall, functionally assigned protein-coding regions have fewer similar copies in the genome, as shown by a significant negative correlation and Gini variable importance above the neutral predictor (Figures 1 & 2). Neither the functionally assigned short or long ncRNAs have significantly different copy-numbers to the negative control sequences (Figure 1).

An alternative approach is to identify repeat-free regions of the genome that could be indicative of functional elements, as these regions may be intolerant of insertions, with this feature now being widely used for identifying bacterial genes required for growth (Cain et al. 2020; Simons et al. 2006). The distance-to-repeat for each sequence was determined using two methods, which either measured the minimum or combined distance to the nearest non-overlapping upstream and downstream repetitive element. Both distance-to-repeat features are significantly positively correlated with all functionally assigned sequence types, with the protein-coding sum of distance-to-repeats and lncRNA minimum distance-to-repeat being the only features to have a Gini variable importance below the neutral predictor (Figures 1 & 2). However, since the protein-coding and lncRNA distance-to repeat features are ranked near the neutral predictor, this suggests that these features capture less information about functionality overall for these sequence types, compared to the other features used by the model (Figure 2).

#### Protein-coding potential features

A number of methods have been developed for differentiating between coding and non-coding sequences, using metrics such as open-reading frame length, codon usage bias and weighted nucleotide frequencies (Lin et al. 2011; Kang et al. 2017; Wang et al. 2013; Fickett 1982). While it is unlikely that coding potential can be used to differentiate between ncRNAs and negative control regions, it is likely that assigned functional protein-coding genes can be classed as “coding” whereas the negative controls will be “non-coding”. Two metrics are used to capture coding potential information, with RNAcode using multiple sequence alignments to calculate a normalised substitution score, whereas the Fickett score uses compositional bias between codon positions from individual sequences (Washietl et al. 2011). The RNAcode score produces a significant positive correlation for protein-coding nucleotide alignments, whilst the short ncRNAs and lncRNAs produce negative correlations, as expected given that this is a measure of protein-coding potential (Figure 1). Since only the protein-coding RNAcode score ranks above the neutral predictor by the random forest, this suggests that the RNAcode score is likely associated with protein-coding functionality (Figure 2). Both the protein-coding and lncRNA Fickett scores produce significant positive correlations with assigned functionality and also rank above the neutral predictor (Figures 1 & 2). However, given that this feature is for detecting coding potential, it is unlikely that the lncRNA Fickett score is producing a meaningful signal and therefore cannot be concluded to be associated with lncRNA functionality.

#### RNA structure features

Functional ncRNAs, and some protein-coding messenger RNAs (mRNAs), are likely to have conserved RNA secondary structures which can be detected with pairwise covariation statistics from sequence alignments (Kertesz et al. 2010; Li et al. 2012; Ding et al. 2014; Rivas et al. 2017; Chiu and Kolodziejczak 1991). Our metrics for capturing the covarying sites in alignment columns, which indicate conserved RNA base-pairs, are the maximum G-test covariance score and minimum covariance E-value produced based on simulated alignments (Rivas et al. 2017). Since functional ncRNAs are also likely to form secondary structures with a lower free energy, we measured the minimum free energy (MFE) structure using either a singular sequence or sequence alignment (Lorenz et al. 2011). Accessibility, which is a measure of the availability of nucleotides for base-pairing, has also been shown to be important for mRNA translation rates, and thus could be associated with functionality (Bhandari et al. 2019). Both covariation features produce significant correlations for all functionally assigned sequences, with short ncRNA minimum covariance E-value and maximum covariance for all sequence types ranking above the neutral predictor, making the latter highly associated with functionality (Figures 1 & 2). Both the protein-coding and lncRNA RNAalifold score, which represents the sequence alignment MFE, produce significant correlations to assigned functionality, with the protein-coding score additionally ranking above the neutral predictor (Figures 1 & 2). Single sequence MFE and accessibility produce significant correlations to assigned short ncRNA functionality and both rank above the neutral predictor (Figures 1 & 2).

#### RNA:RNA interactions features

While RNA interactions are important for regulation, the opposite may also be true, with evidence suggesting that selection may occur against stochastic interactions for different ncRNAs, resulting in RNA:RNA interaction avoidance signals (Mann et al. 2017; Umu et al. 2016). We approximated the number of RNA:RNA interactions by computing the minimum and average interaction energies between each sequence and a selection of 34 highly-expressed genes. Both interaction features are significantly correlated and rank above the neutral predictor by the random forest for short ncRNAs (Figures 1 & 2). In contrast, the minimum interaction energy is significantly and inversely correlated for the protein-coding sequences but ranks below the neutral predictor (Figures 1 & 2). For minimum interaction energy, short ncRNAs have a positive correlation of 0.27 which suggests that excess RNA:RNA interactions are avoided, whereas a negative correlation of −0.10 for protein-coding exons implies there are more RNA:RNA interactions than expected (Figures S1 & S2). Since only protein-coding sequences have a significantly negative correlation between the minimum interaction energy and accessibility (Figures S1 & S2), then a difference in sequence accessibility could be driving these observations, rather than the interactions observed (Busch et al. 2008).

#### Population variation features

The amount of variation between individuals in a population is likely to be influenced by functionality, with excess variation in functionally important regions being selected against and little-to-no selection acting on non-functional regions. To assess this, we used 1,000 Genomes Project (1kGP) and Genome Aggregation Database (gnomAD) data to calculate SNP count per region, SNP density per region and the average minor allele frequency (MAF) observed (The 1000 Genomes Project Consortium 2015; Karczewski et al. 2020). A high SNP count and density implies there is excess variation, which is more likely to occur in non-functional regions, whereas a lower MAF may result from the more common allele being more conserved across populations, and hence more likely to be functional. Both SNP count and density derived from 1kGP and gnomAD data are significantly correlated with assigned short ncRNA functionality, with there being no significant correlation between the population features and assigned lncRNA functionality (Figure 1). All gnomAD derived features and 1kGP average MAF are significantly correlated with assigned protein-coding functionality (Figure 1). Of these significantly correlated features, only 1kGP SNP density ranks above the neutral predictor for short ncRNAs, with all remaining correlated population features ranking below the neutral predictor (Figure 2).

Since both short ncRNA SNP count and density have a positive correlation of 0.28 to assigned functionality, this implies that the short ncRNAs contain more genetic variation than matched control regions, which is opposite to the expected negative correlation between SNP features and sequence conservation, as observed for protein-coding regions (Figure S1 & S2). To highlight the issue within the SNP data, functionally assigned short ncRNAs with high SNP densities were compared to protein-coding and lncRNA exons with the same sequence length, with high SNP density being defined as greater than both the median gnomAD and 1kGP SNP density for each sequence type (Figure S5A & S5B). The short ncRNAs show three clear peaks at sequence lengths 72, 73 and 82 bp, meaning they have greater SNP densities than protein-coding or lncRNA exons of the same length, with the majority of these short ncRNAs being tRNAs (Figure 3). However, it may be possible the correlations observed are caused by the short ncRNA negative controls, since there is a slight significant difference between the SNP densities for short ncRNA and protein-coding negative control sequences, with the latter performing as expected (Figure S5C & S5D).

**Figure 3:**
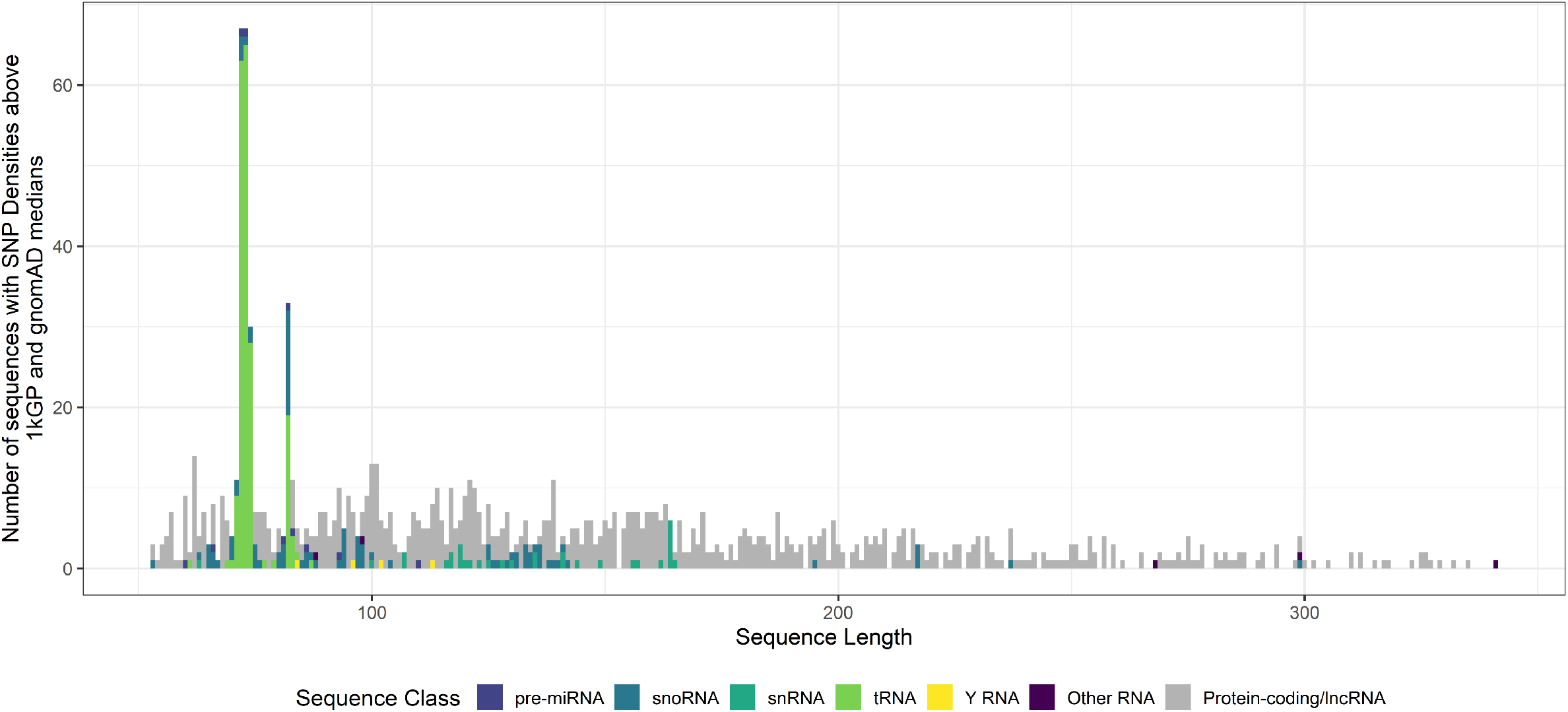
Bar graph of assigned functional sequences with high SNP densities. A Bar graph was used to highlight the number of assigned functional sequences with high SNP densities, which are defined as sequences with SNP densities above the median for both gnomAD and 1kGP (Figure S5). High SNP density protein-coding and lncRNA sequences are shown in grey to show the “expected” SNP density for sequences of the same length. The “Other RNA” class contains Vault RNA, RNA Component of Mitochondrial RNA Processing Endoribonuclease, Ribonuclease P RNA and signal recognition particle RNA. Sample sizes are as follows: pre-miRNA (N=10), snoRNA (N=89), snRNA (N=32), tRNA (N=195), Y RNA (N=4), Other RNA (N=5), Protein-coding (N=426) and lncRNA (N=481).

## Discussion

The two interpretations of “function” in genetics that are broadly used either require only biochemical activity, such as transcription, or evidence of both biochemical activity and evolutionary selection (Eddy 2012; Doolittle and Brunet 2017; Graur 2017). In this study, we find a total of 19 short ncRNA, 17 protein-coding and nine lncRNA features associated with the functionality of each respective sequence type (Figures 1 & 2; Table S3). Sequence conservation features were the most strongly associated with protein-coding sequence functionality, while evidence of transcription was the most associated with ncRNA functionality (Figures 1 & 2). Features that were associated with functionality, regardless of sequence type, were also from these two groups of features with the exception of maximum covariance, which suggests that a combination of biochemical activity and evolutionary selection would produce a more robust definition of function (Figure 2; Table S3). While transcription is also associated with assigned lncRNA functionality, there is no guarantee that these are informative features due to the low performing lncRNA classification models, reflecting the underlying complexity of lncRNAs (Table 1 & S3).

Our results indicated that functionally assigned short ncRNAs tend to have an excess of SNPs, with short ncRNA SNP densities on average being 3.37 and 4.39 times larger than their matched negative controls for gnomAD and 1kGP respectively (Figures 3 & S5). While this issue appears to primarily affect tRNAs (Figure 3), an excess of SNPs raises concerns about genome variation data since many short ncRNAs are among the most conserved genes in all life (Hoeppner et al. 2012). Previous studies have identified issues with entire databases or studies such as the 1kGP, with the main cause of false positive SNPs being the reference genome quality due to misassembled, absent regions or collapsed duplications in the genome, which contaminate SNP data (Ribeiro et al. 2015; Hartasánchez et al. 2018; Guo et al. 2017; Anderson-Trocmé et al. 2020; Mafessoni et al. 2018). Considering that the high SNP density observed in tRNAs was also present in the more recent and larger gnomAD database (Figure 3), then it is likely that duplicates of the same tRNA, pseudogene copies or tRNA derived repeats have been mapped to a single set of chromosomal coordinates, thus inflating the number of SNPs present (Bermudez-Santana et al. 2010; Frenkel et al. 2004; Karczewski et al. 2020). These possibilities suggest that collapsed duplications and misassembled genomic regions still contaminate modern SNP studies, and until such artefacts are resolved, conclusions drawn from SNP data should be regarded carefully (Anderson-Trocmé et al. 2020).

Maximum covariance stands out as being highly predictive of all functionally assigned sequences (Table S3), which is unusual for the protein-coding sequences given that this feature was originally designed for conserved RNA secondary structure analysis (Rivas et al. 2017; Rivas 2020). While mRNA secondary structure has long been proposed to be important for regulating and controlling translation rates (Pelletier and Sonenberg 1987; de Sousa Abreu et al. 2009), the covariation signatures we observe in protein-coding nucleotide alignments are unlikely to be due to conserved secondary structure since the full-length mRNA was not analysed. Instead, this may be due to compensatory variation between codon positions one and two, with this signal in mRNAs having been previously reported (Rivas 2020). We verified this by analysing the distribution of distances between covarying alignment positions for the maximum covariance recorded per sequence (Figure S6). The most frequent distance between covarying alignment columns is one, with there being modes at factors of three from distances six up to thirty (Figure S6). This periodicity of three nucleotide distances between modes in the distribution further supports the signal being from between and within codon compensation, rather than from RNA structure (Figure S6).

In comparison to short ncRNA and protein-coding sequences, lncRNAs have the least defined functional signals, as evident by weak correlations between features and assigned lncRNA functionality and relatively poor performing classification models (Figure S3; Table 1). Previous studies investigating lncRNA functionality have reported AUC values between 0.550 and 0.990, compared to 0.722 to 0.835 (Table 1), and sensitivity values ranging from 0.404 to 0.619, compared to 0.374 to 0.410 (Table 1), for their classification models (Tsai et al. 2017; Simopoulos et al. 2018). However, these differences in model performance are likely due to method variation, such as scoring lncRNAs using a protein-coding trained model, rather than a lncRNA trained model, or using protein-coding or pseudogenes as negative controls, rather than unannotated sequences (Tsai et al. 2017; Simopoulos et al. 2018). A common theme between these studies and ours is the incorporation of numerous features to most accurately determine functionality (Table S3), implying that multiple criteria are required to form a robust definition for functionality (Simopoulos et al. 2018; Tsai et al. 2017). While lncRNA annotation methods tend to adopt a similar approach, the less defined lncRNA functionality signal suggests that non-genic sequences are being annotated as lncRNAs, meaning current thresholds may not be accounting for experimental and biological noise (Xu et al. 2017).

While we find that sequence conservation and transcriptome expression are most strongly associated with assigned protein-coding and ncRNA functionality respectively, limitations in our method design may have also contributed to these differences (Figures 1 & 2). Despite analysing a total of 211 tissue and primary cell RNA-Seq datasets, it is still possible that sequences that had no transcription recorded could be a result of cell or condition specific transcription that were not present in the datasets analysed. PhyloP and PhastCons sequence conservation scores, which were the highest performing short ncRNA and protein-coding conservation features (Figure 1), are derived from dated sequence assemblies (2005-2013) and could contain inaccurate sequences or alignments (Kitts et al. 2016; Haeussler et al. 2019). GERP scores, which should have benefitted from the use of improved sequence alignment tools and incorporating recently released genomes (Cooper et al. 2005), tend to be less associated with functionality (Figure 1). While it may be possible that our negative control sequences are unsuitable for this study, more ideal controls are not currently widely available. The overall effect of our results shows that by considering these features together when investigating functionality, thus creating more stringent criteria, we can ensure more genes and non-genic regions are correctly classified.

## Materials and Methods

In the following we will describe the positive and negative control annotations we selected, the genome features we used for the analysis and the analytical methods we used to identify features associated with functionality.

### Retrieval of functional genes

An initial dataset of 19,031 protein-coding sequence IDs was obtained from HGNC, which were further filtered to exclude those with withdrawn entries and had multiple RefSeq IDs associated with one gene (Braschi et al. 2019). RefSeq IDs were used to obtain the chromosome coordinates and corresponding sequences from the GRCh38.p13 human genome GFF file (RefSeq assembly accession: GCF_000001405.39) (O’Leary et al. 2016). Functional short ncRNAs and multi-exonic lncRNAs were obtained from RNAcentral v15, which were filtered to include those in the HGNC database, giving an initial total of 959 short ncRNAs and 3,959 lncRNAs (The RNAcentral Consortium 2019). An additional 1,862 precursor miRNAs were obtained separately to the short ncRNA dataset, as they are overrepresented within RNAcentral and have unusually stable structures (Freyhult et al. 2005; The RNAcentral Consortium 2019). Precursor miRNAs were selected at random to bring the total short ncRNA dataset to 1,000.

### Negative control sequences

Negative control sequences were sampled from the GRCh38.p13 human genome (O’Leary et al. 2016), and then filtered to exclude sequences with undetermined nucleotides (i.e. “N”). For consistency, the length of the negative control regions was based on the corresponding functional sequences. Bedtools 2.29.0 was used to filter out controls that overlapped with Swiss-Prot or GENCODE annotations (Quinlan 2014; Haeussler et al. 2019; Harrow et al. 2012). Post-filtering, there were 895 short ncRNA negative controls, 790 lncRNA negative controls and 1,104 protein-coding negative controls.

### Feature inclusion criteria

In order for a potential feature of function to be included in the analysis, it needed to meet the following criteria:

1. A genome-wide statistic (ie: applicable to negative control sequences).
2. Informative and non-redundant with other included features.
3. Unbiased towards known genes (e.g. not based upon homology to known genes).
4. Readily accessible for the GRCh38 genome.
5. Have a reasonably installable package.

From our initial selection of 52 features, 23 were removed as they did not meet our inclusion criteria, and 29 features were kept for further analysis (Table S1).

### Intrinsic sequence features

The main intrinsic sequence feature analysed was G+C content, which was calculated by finding the percentage of guanines and cytosines per sequence. To identify features that were performing worse than random, the 5’ chromosomal coordinate for each sequence was also included as a “feature”, since it is unlikely to significantly contribute to functionality.

### Sequence conservation features

The average and maximum PhyloP and PhastCons scores used to analyse sequence conservation were extracted from the UCSC hg38 phastCons100way and phyloP100way bigwig files using UCSC genome browser ‘kent’ bioinformatic utilities v385 (Haeussler et al. 2019). The average and maximum GERP scores for each region were extracted using Bedtools 2.29.0 from a release 101 Ensembl multiple genome alignment using 111 mammals (Cooper et al. 2005; Yates et al. 2020; Quinlan 2014).

### Transcriptome expression features

To measure the level of potential transcription in each sequence, total and small RNA-Seq datasets from 71 human tissue and 140 human primary-cell samples were obtained from ENCODE (ENCODE Project Consortium 2012). Both datasets have rRNAs depleted, with the total RNA-Seq not including transcripts shorter than 200 bp long and small RNA-Seq only including transcripts less than 200 bp long (ENCODE Project Consortium 2012). Lists of the ENCODE dataset IDs used are available on Github. Samtools 1.10 was used to determine the MRD and RPKM for each RNA-Seq dataset, with the maximum across all datasets being recorded for each sequence (Li et al. 2009).

### Genomic repeat associated features

To estimate the genomic copy number of each sequence, BLAST 2.10.0 was used to identify all possible alignments against version GRCh38.p13 of the human genome (O’Leary et al. 2016; Altschul et al. 1990). Distance-to-repeat features were calculated using non-redundant hits of repetitive DNA elements in the human genome obtained from Dfam v3.1, and the closest non-overlapping repetitive elements up and downstream of the sequence of interest were identified with Bedtools 2.29.0 (Quinlan 2014; Hubley et al. 2016). Both the minimum and the sum of the distances to the nearest up and downstream repeats were recorded.

### Protein and RNA specific features

#### Coding potential

The coding potential of each sequence was calculated using CPC2 beta and RNAcode (ViennaRNA 2.4.14), which were either based on Fickett scores from individual sequences, or RNAcode scores from sequence alignments (Kang et al. 2017; Lorenz et al. 2011).

#### RNA structure

Multiple sequence alignments were obtained from the UCSC multiz100way alignment using the UCSC genome browser ‘kent’ bioinformatic utilities v385, and the consensus secondary structures were generated using RNAalifold (ViennaRNA 2.4.14) (Haeussler et al. 2019; Lorenz et al. 2011). R-scape 1.4.0 was used to calculate maximum covariance score and minimum covariance E-value (Rivas et al. 2017). RNAfold and RNAplfold from ViennaRNA 2.4.14 were used to calculate both MFE and accessibility values for each region (Lorenz et al. 2011; Bhandari et al. 2019).

#### RNA:RNA interactions

To determine whether RNA interactions or avoidance are associated with sequence functionality, IntaRNA 3.2.0 was used to determine minimum and average interaction free energies between control sequences and a curated database of 34 human ncRNA sequences (Table S4) from RNAcentral v15 (The RNAcentral Consortium 2019). These 34 ncRNAs are known to be abundant and interact with a variety of RNAs (Driedonks and Nolte-’t Hoen 2018; Marvin et al. 2011; Yan et al. 2019; Mann et al. 2017).

### Population variation features

To explore population variation, we used Tabix 1.10 to obtain data from phase 3 of the 1kGP and gnomAD v3 (The 1000 Genomes Project Consortium 2015; Li 2011; Karczewski et al. 2020). When obtaining the 1kGP data, chromosome coordinates were lifted over from the GRCh38 to the GRCh37 reference genome using the UCSC genome browser ‘kent’ bioinformatic utilities v385 (Haeussler et al. 2019). Both the SNP count and density were recorded, with density being calculated by dividing the SNP count by sequence length. The 1kGP average MAF for each sequence was obtained using VCFtools 4.2 (Danecek et al. 2011).

### Analysis of potential functionality features

Spearman correlation matrices were generated in R 3.6.0, with the confidence intervals calculated using a bootstrap approach (N=1,000). Random forests were generated in R with 1,000 trees, 70% training data and 30% test data using randomForest v4.6-14 (Liaw et al. 2002). A bootstrap approach was used to simulate a further 100 classification models to assess performance variation due to sampling biases.

### Data Access

All data described and downloaded were last accessed in September 2020. All custom scripts, information on downloaded primary data and the generated datasets are available on GitHub: https://github.com/helena-bethany/gene-functionality

## Acknowledgements

This project has benefitted from discussions and advice from Eric Nawrocki. Assistance with identifying issues with SNP data was given by Moizle Ocariza and Harry Biggs. We thank Elena Rivas, Austen Ganley, Alex Gavryushkin and Sarah Diermeier for comments on the draft manuscript and Bikash Kumar Bhandari for assisting in computing RNA structure avoidance values. HBC receives funding from the University of Otago Summer Research Scholarship. PPG receives funding from the 2019 Endeavour Fund (Smart Ideas) and the 2019 Strategic Science Investment Fund (Data Science Research Programme) from the Ministry of Business, Innovation and Employment, New Zealand.

