## Supplemental_Figures for "Features of Functional Human Genes"

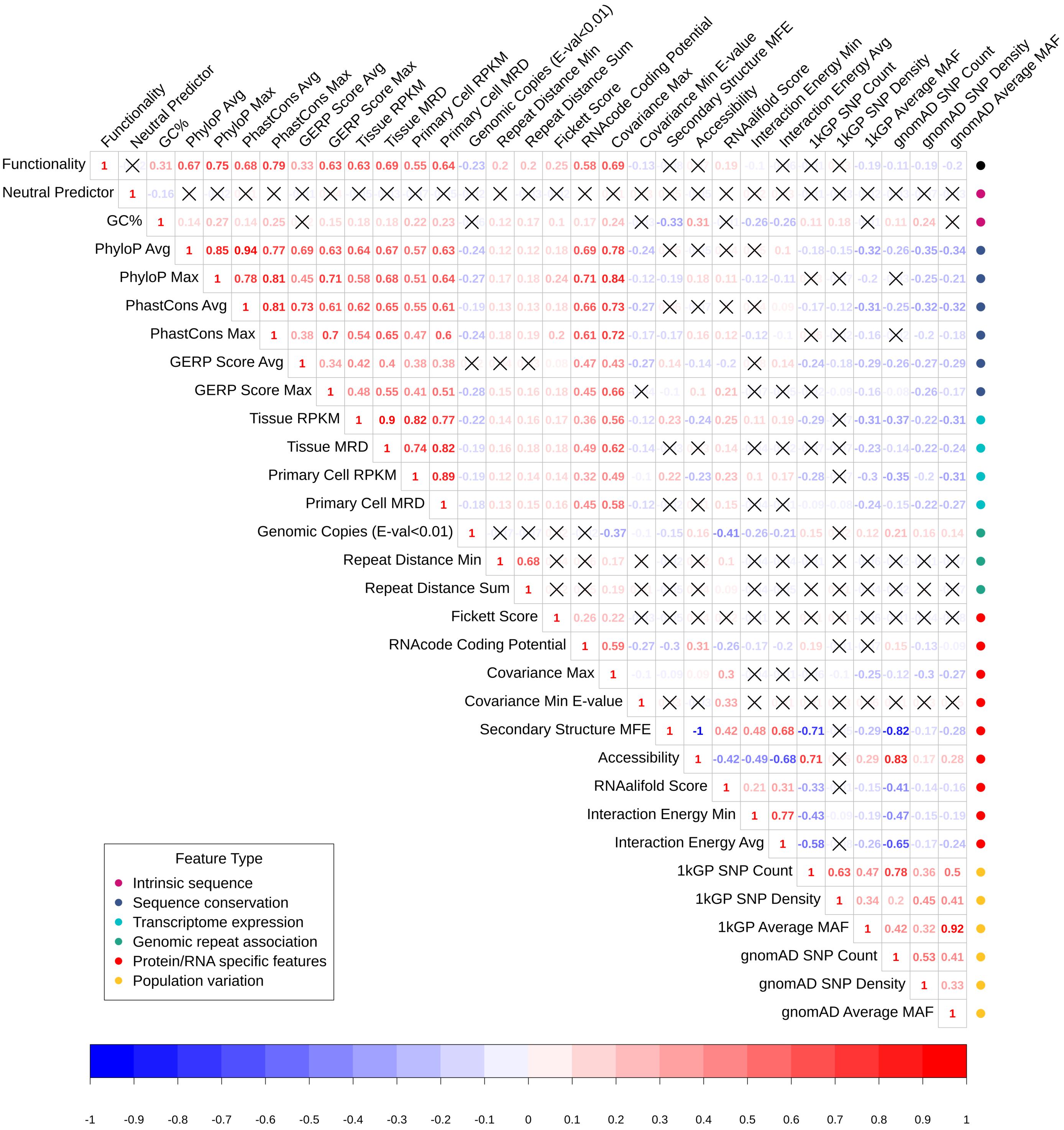


**Figure S1: Protein-coding exon correlation matrix.** Spearman correlation of functionality predictors for protein-coding sequences ($N^{Functional}$=1,576 & $N^{Non\text{-}functional}$=845). The top row of the matrix shows the correlation of each feature with the functional category, with positively correlated features in red and negatively correlated features in blue. Spearman's Rho was used to determine the significance of each correlation, which were corrected for multiple testing using a Bonferroni correction. Non-significant correlations with a p-value > 0.05 have been marked with a cross.

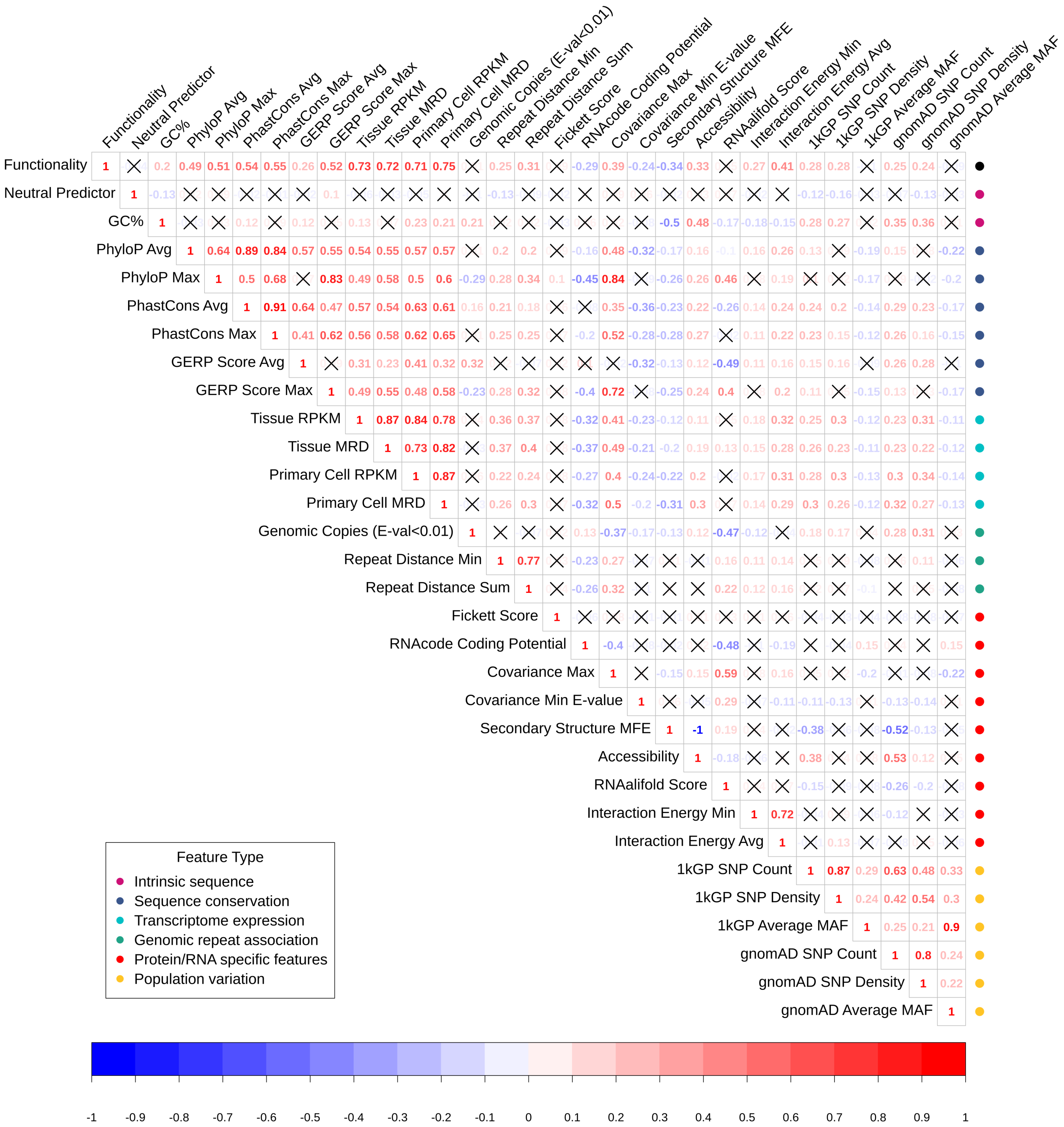


**Figure S2: Short ncRNA correlation matrix.** Spearman correlation of functionality predictors short ncRNA sequences ($N^{Functional}$=803 & $N^{Non\text{-}functional}$=635). The top row of the matrix shows the correlation of each feature with the functional category, with positively correlated features in red and negatively correlated features in blue. Spearman's Rho was used to determine the significance of each correlation, which were corrected for multiple testing using a Bonferroni correction. Non-significant correlations with a p-value > 0.05 have been marked with a cross.

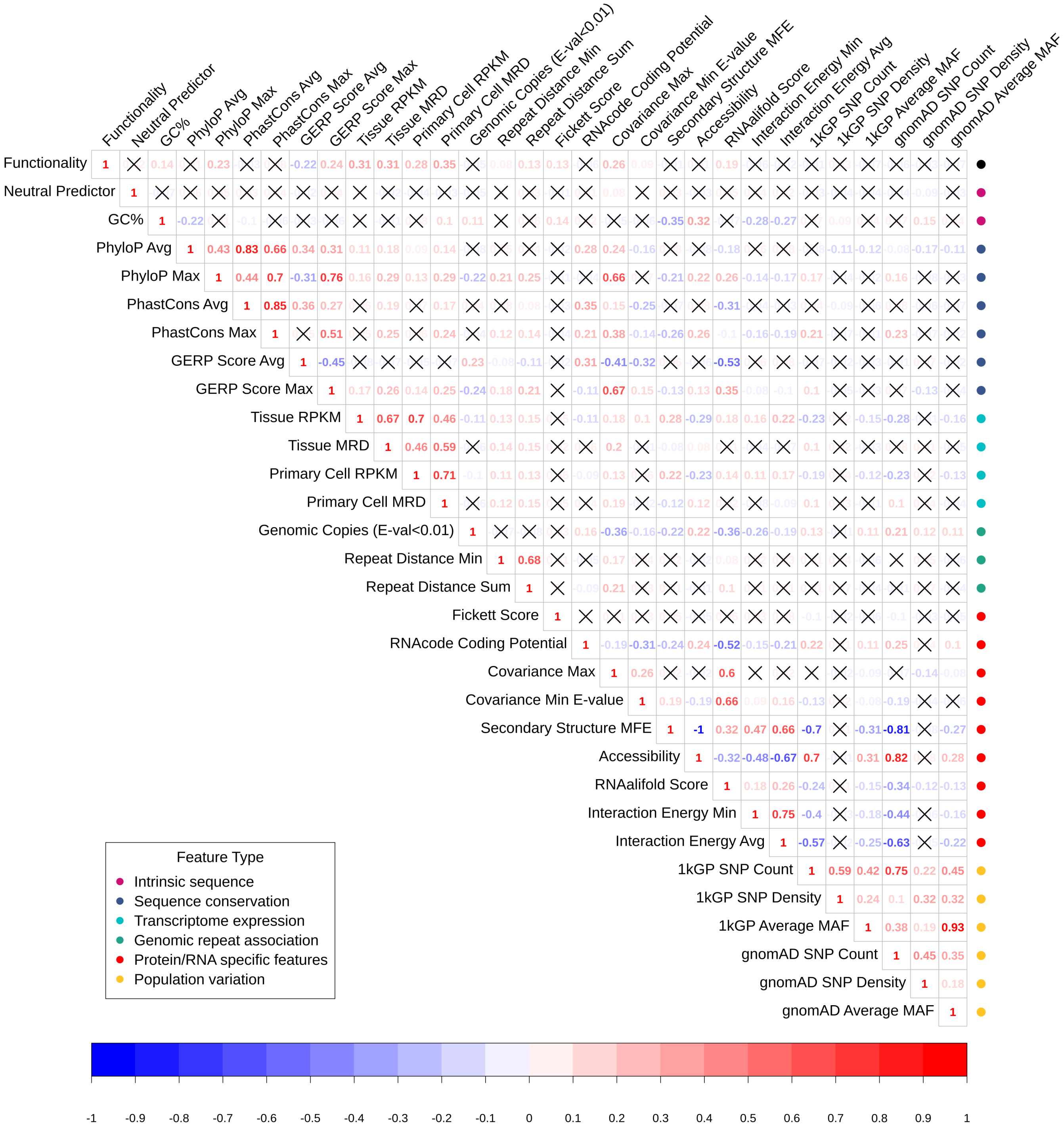


**Figure S3: Long ncRNA correlation matrix.** Spearman correlation of functionality predictors lncRNA sequences ($N^{Functional}$=1,693 & $N^{Non-functional}$=635). The top row of the matrix shows the correlation of each feature with the functional category, with positively correlated features in red and negatively correlated features in blue. Spearman's Rho was used to determine the significance of each correlation, which were corrected for multiple testing using a Bonferroni correction. Non-significant correlations with a p-value > 0.05 have been marked with a cross.

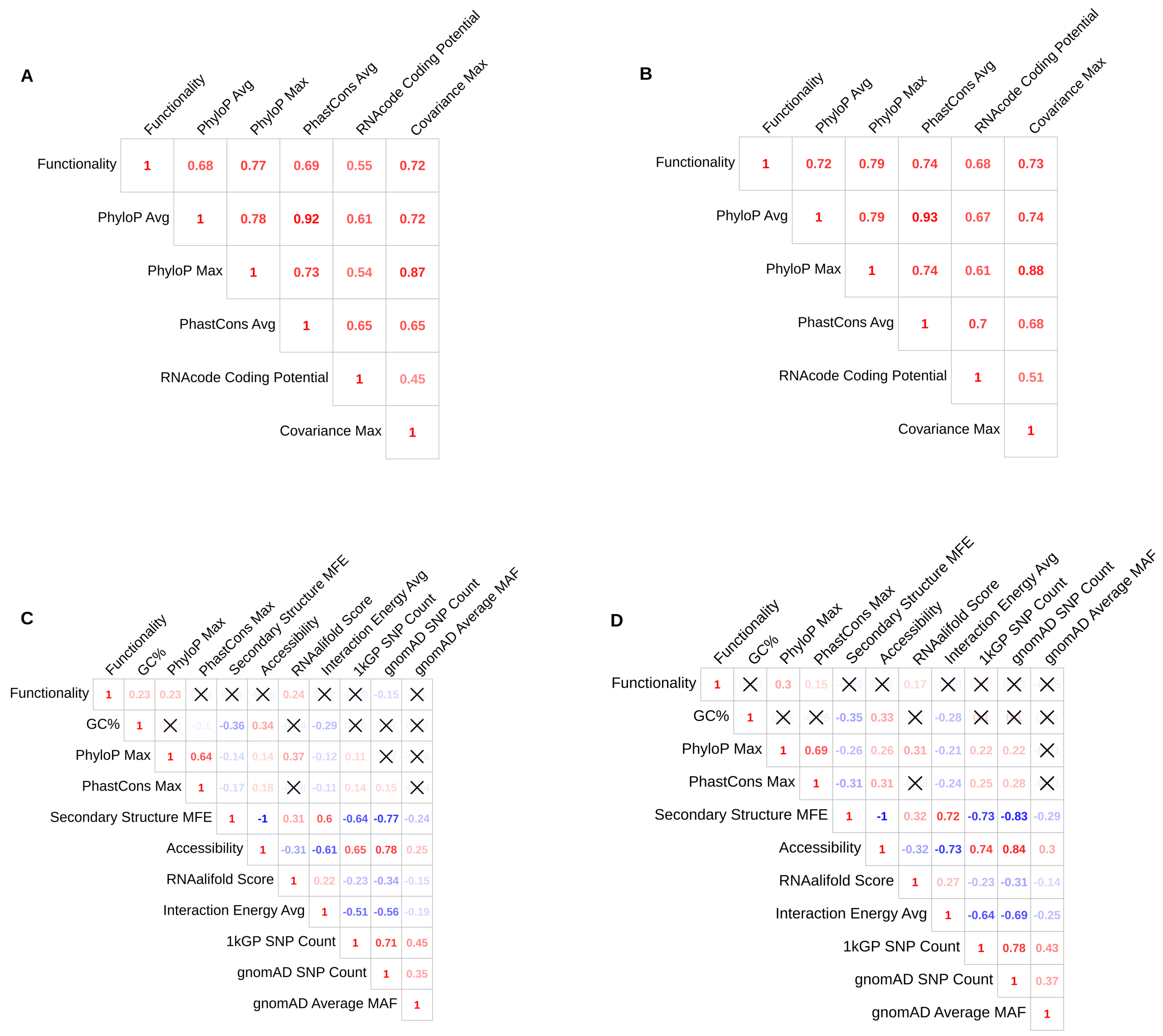


**Figure S4: Protein-coding and lncRNA exon two and three correlation matrices.** Spearman correlation of functionality predictors that have significantly different distributions between either protein-coding or lncRNA exons two and three (Table S2). If a feature does not show the same correlation trends between exons, then it is unlikely that the feature will be representative of the full-length sequence, and thus unlikely to be associated with functionality. The top row of the matrix shows the correlation of each feature with sequence functional, with positively correlated features in red and negatively correlated features in blue. Spearman's Rho was used to determine the significance of each correlation, which were corrected for multiple testing using a Bonferroni correction. Non-significant correlations with a p-value > 0.05 have been marked with a cross. **A:** Protein-coding Exon Two ($N^{Functional}$=797, $N^{Non\text{-}functional}$=845). **B:** Protein-coding Exon Three ($N^{Functional}$=779, $N^{Non\text{-}functional}$=845). **C:** lncRNA Exon Two ($N^{Functional}$=886, $N^{Non\text{-}functional}$=635). **D:** lncRNA Exon Three ($N^{Functional}$=827, $N^{Non\text{-}functional}$=635).

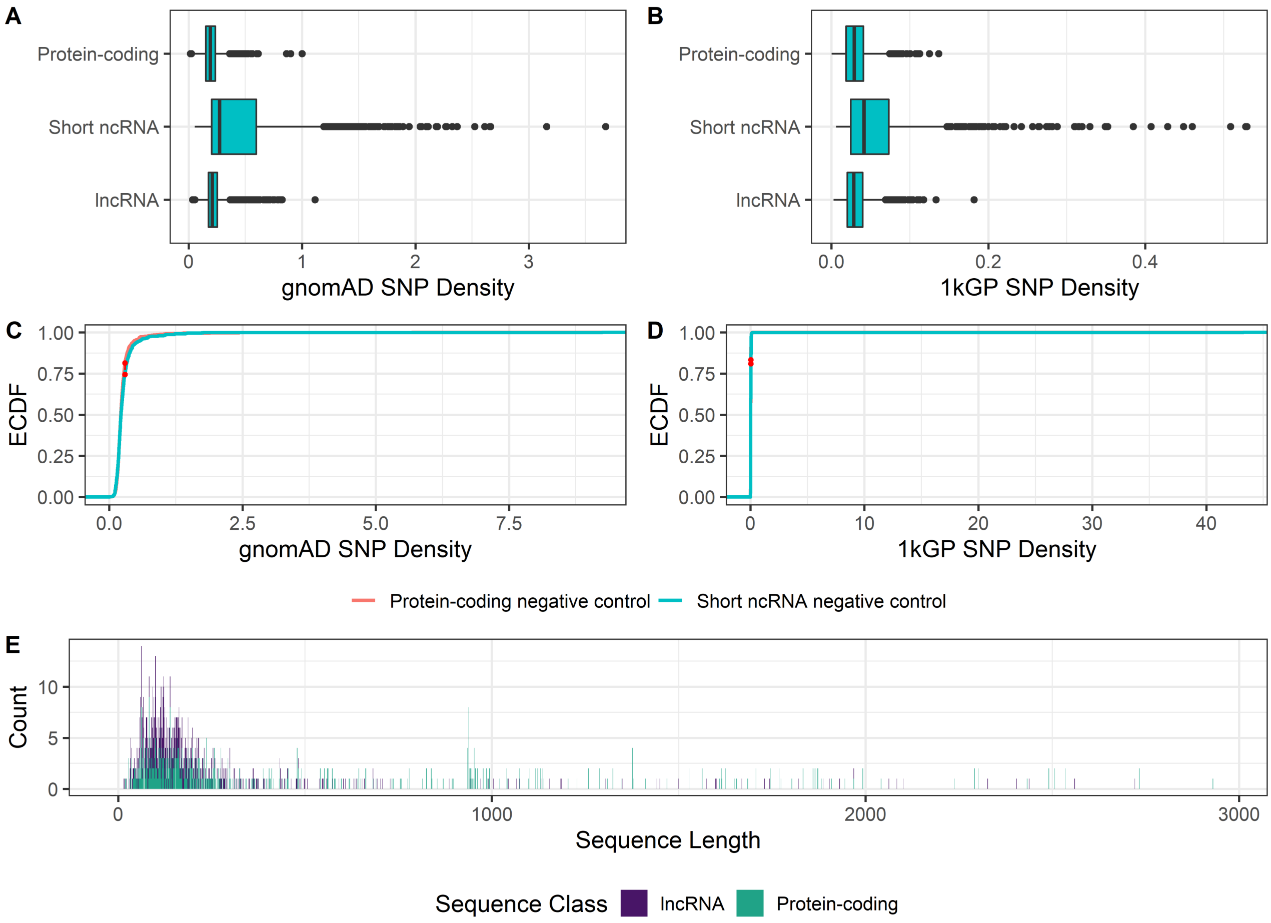


**Figure S5: Comparison of 1kGP and gnomAD SNP densities.** Boxplots were used to identify the threshold for outlier SNP densities in assigned functional sequences, which is any sequence with a SNP density with a value 1.5 times the interquartile range. The Empirical Cumulative Distribution Function (ECDF) was used to visualise the distributions for the negative control sequence SNP Density values, with the significance of the maximum distance (D) between the two distributions (red dotted line) determined using a KS-test. **A:** Protein-coding (Median=0.192, N=1,987), short ncRNA (Median=0.274, N=991) and lncRNA (Median=0.209, N=1,984). **B:** Protein-coding (Median=0.029, N=1,805), Short ncRNA (Median=0.042, N=919) and lncRNA (Median=0.029, N=1,847). **C:** Comparison of gnomAD SNP Densities for short ncRNA (N=878) and protein-coding negative control sequences (N=1,099). D= 0.073 & P-value = 0.012. **D:** Comparison of 1kGP SNP Densities for short ncRNA (N=766) and protein-coding negative control sequences (N=1,020). D= 0.102 & P-value = 2.25e-04. **E:** All protein-coding (N=654) and lncRNA (N=593) exons with gnomAD and 1kGP SNP densities greater than both their respective medians.

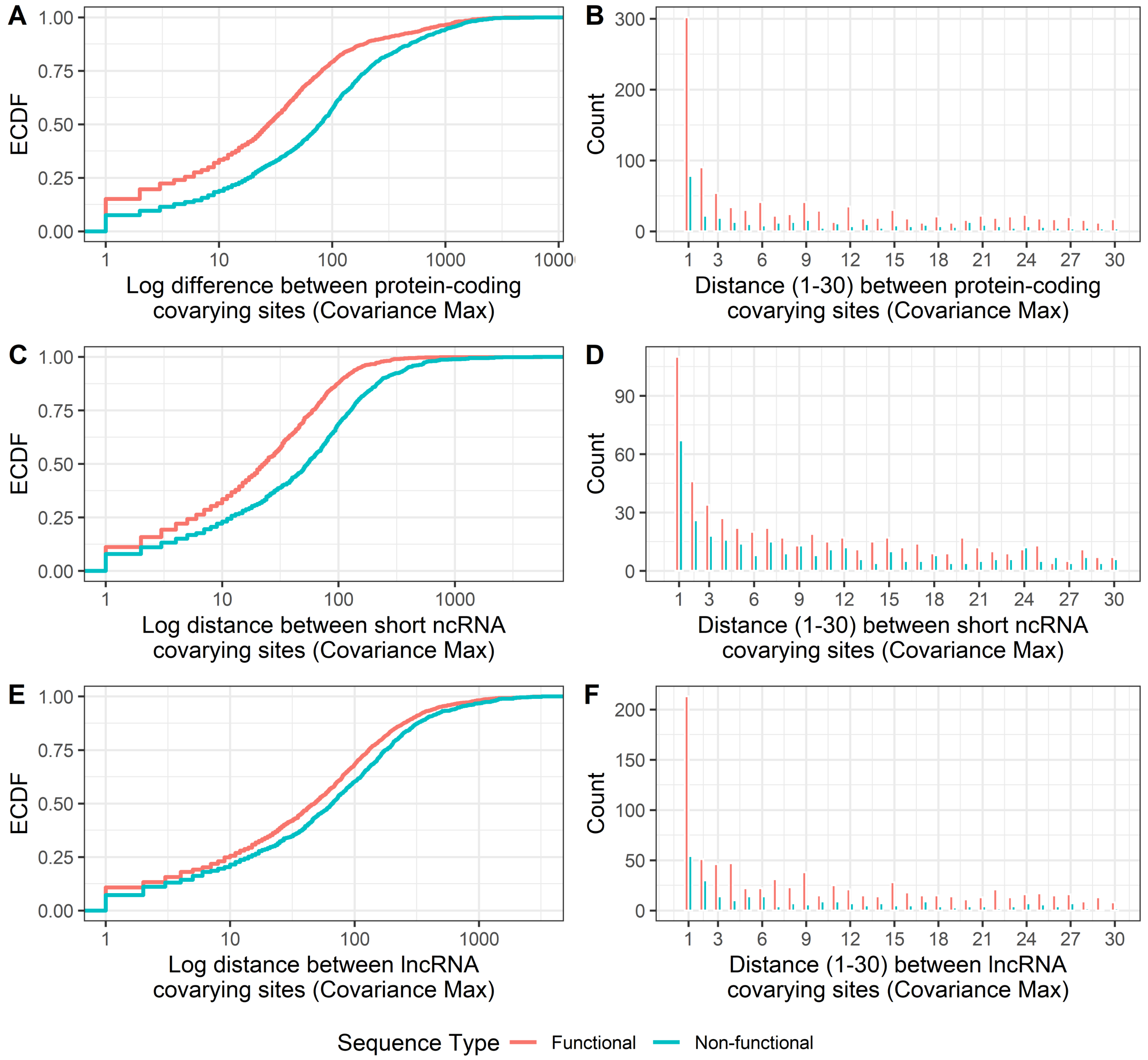


**Figure S6: Distances between covarying sites for functional and negative control sequences.** Distances observed between covarying sites which recorded the maximum covariance for each sequence. Distance was calculated from R-scape data by subtracting the 5` covarying position (left_pos) from the 3` covarying position (right_pos). Panels A-C visualise the ECDF to compare the log difference in covarying site distance between functional (red) and the negative control "non-functional" (blue) sequences. Panels D-F visualise covarying distances ranging between 1 and 30, with functional (red) and negative control (blue) sequences split for each recorded distance. **A:** Protein-coding exons ($N^{Exon2}$=999, $N^{Exon3}$=997, $N^{Non\text{-}functional}$=1038). **B:** Short ncRNAs ($N^{Functional}$=983, $N^{Non\text{-}functional}$=841). **C:** lncRNAs exons ($N^{Exon2}$=990, $N^{Exon3}$=990, $N^{Non\text{-}functional}$=753). **D:** Protein-coding exons ($N^{Exon2}$=507, $N^{Exon3}$=540, $N^{Non\text{-}functional}$=336). **E:** Short ncRNAs ($N^{Functional}$=555, $N^{Non\text{-}functional}$=325). **F:** lncRNAs exons ($N^{Exon2}$=412, $N^{Exon3}$=413, $N^{Non\text{-}functional}$=259).
