## Supplemental_Tables_S2_S4 for "Features of Functional Human Genes"

**Table S2: Inter-exon differences.** P-values from two sample KS-tests comparing the distribution between exons two and three of either the protein-coding or lncRNA sequence. These were corrected for multiple testing using a Bonferroni correction (N=30).

| | Protein-coding | lncRNA |
|---|---|---|
| Neutral Predictor | 1.00 | 1.00 |
| GC% | 1.00 | **3.22e-07** |
| PhyloP Max | **1.08e-04** | **8.76e-04** |
| PhyloP Avg | **1.20e-03** | 1.00 |
| PhastCons Max | 1.00 | **5.79e-04** |
| PhastCons Avg | **0.037** | 0.105 |
| GERP Score Max | 1.00 | 0.407 |
| GERP Score Avg | 0.713 | 1.00 |
| Tissue RPKM | 1.00 | 0.336 |
| Tissue MRD | 1.00 | 1.00 |
| Primary Cell RPKM | 1.00 | 1.00 |
| Primary Cell MRD | 1.00 | 1.00 |
| Genomic copies (E-val<0.01) | 1.00 | 1.00 |
| Repeat Distance Min | 1.00 | 1.00 |
| Repeat Distance Sum | 0.388 | 1.00 |
| Fickett Score | 1.00 | 0.137 |
| RNAcode Coding Potential | **7.21e-05** | 1.00 |
| Covariance Max | **4.19e-03** | 1.00 |
| Covariance Min E-value | 1.00 | 0.110 |
| Secondary Structure MFE | 1.00 | **1.60e-08** |
| Accessibility | 1.00 | **3.75e-07** |
| RNAalifold Score | 1.00 | **0.044** |
| Interaction Energy Min | 1.00 | 0.251 |
| Interaction Energy Avg | 1.00 | **4.44e-08** |
| 1kGP SNP Count | 1.00 | **3.50e-10** |
| 1kGP SNP Density | 1.00 | 1.00 |
| 1kGP Average MAF | 1.00 | 0.356 |
| gnomAD SNP Count | 1.00 | **6.60e-15** |
| gnomAD SNP Density | 1.00 | 0.743 |
| gnomAD Average MAF | 1.00 | **0.049** |

**Table S4:** Summary of the 34 RNA sequences in the curated RNA:RNA interaction database.

| RNA Type | Total | RNAcentral IDs |
|---|---|---|
| tRNA | 20 | URS00003D2CC9_9606 (Ala), URS000038803E_9606 (Val), URS0000735371_9606 (Leu), URS000072E1AF_9606 (Ile), URS00006A14B6_9606 (Cys), URS000065DEBF_9606 (Met), URS0000C8E9D4_9606 (Phe), URS0000233681_9606 (Tyr), URS000047B05D_9606 (Thr), URS00001A72CE_9606 (Trp), URS0000639DBE_9606 (Glu), URS00006D74B2_9606 (Asp), URS0000659172_9606 (Asn), URS000074C9DF_9606 (Gln), URS0000733374_9606 (Ser), URS000067A424_9606 (His), URS000007E37F_9606 (Arg), URS000034AAC2_9606 (Gly), URS0000744456_9606 (Pro), URS0000C8E9CE_9606 (Lys) |
| 5S rRNA | 1 | URS00000F9D45_9606 |
| Spliceosomal RNA | 9 | URS00006BDF17_9606 (U1), URS00003EE995_9606 (U2), URS00003F07BD_9606 (U4), URS000075BAAE_9606 (U5), URS000035C796_9606 (U6), URS000000A142_9606 (U11), URS000075EF5D_9606 (U12), URS000075ADBA_9606 (U4atac), URS0000443498_9606 (U6atac) |
| Y RNA | 3 | URS0000103047_9606 (Y1), URS00005CF03F_9606 (Y3), URS0000007D24_9606 (Y4) |
| Ribonuclease P | 1 | URS000013F331_9606 (H1) |
